# A hybrid combination of in vitro cultured buccal mucosal cells using two different methodologies, complementing each other in successfully repairing a stricture-inflicted human male urethral epithelium

**DOI:** 10.1101/2023.08.29.555240

**Authors:** Akio Horiguchi, Toshihiro Kushibiki, Mayumi Yoshine, Masayuki Shinchi, Kenichiro Ojima, Yusuke Hirano, Shojiro Katoh, Masaru Iwasaki, Vaddi Surya Prakash, Koji Ichiyama, Rajappa Senthilkumar, Senthilkumar Preethy, Samuel JK Abraham

## Abstract

**Background:** Autologous buccal mucosal tissue derived cell transplantation techniques in repairing a stricture inflicted male urethral epithelium have been evolving. There was not much of clarity on the cell type, in vitro culture methods and the mode of transplantation, until we reported our buccal epithelium expanded and encapsulated in scaffold-hybrid approach to urethral stricture (BEES-HAUS) clinical study yielding a successful engraftment and repair with a long-term patency. We herein report with technical clarity on the advantages of mixing two-dimensional (2D) monolayer cultured fibroblast like cells and three dimensional (3D) thermo-reversible gelation polymer (TGP) cultured cells; the former secreting IGF-1, a cytokine known for its healing effects and the latter expressing epithelial surface markers in flow cytometry, both sourced from human buccal tissue, together transplanted using TGP as a carrier.

**Methods:** Human buccal tissues (n=22) redundant after urethroplasty surgery was used after informed consent and IEC approval. They were enzyme digested, divided into two portions; one was cultured as monolayer method (2D) and the other in 3D in TGP. Flowcytometry and quantification of IGF-1 in cell culture supernatant through the culture period were undertaken.

**Results:** In flowcytometry, the cells on day 0, lacked AE1/AE3 - pancytokeratin expression indicative of epithelial phenotype of culture, which progressively increased in the 3D-TGP group, during invitro culture of up to 21 days. The 2D showed expression of only fibroblasts like cells that were negative for AE1/AE3 but positive for CD140b. IGF-1 secretion was significantly higher in 2D cultures than in 3D-TGP (p-value < 0.05).

**Conclusion:** The 3D- TGP cultured cells of epithelial nature and the 2D cultured fibroblast like cells secreting IGF-1, together when transplanted using TGP scaffold as a carrier, adapted to a hostile in vivo milieu after releasing the fibrous strands with urethrotomy, successfully engrafted and repaired a stricture-inflicted male urethral epithelium in the BEES-HAUS procedure. While this hybrid combination of cells are considered to have potential in managing other epithelial damages, further research of such hybrid combination and their behaviour in disease affected environments may help to expand this solution in regenerating and repairing other tissues and organs as well.

**Graphical abstract:** 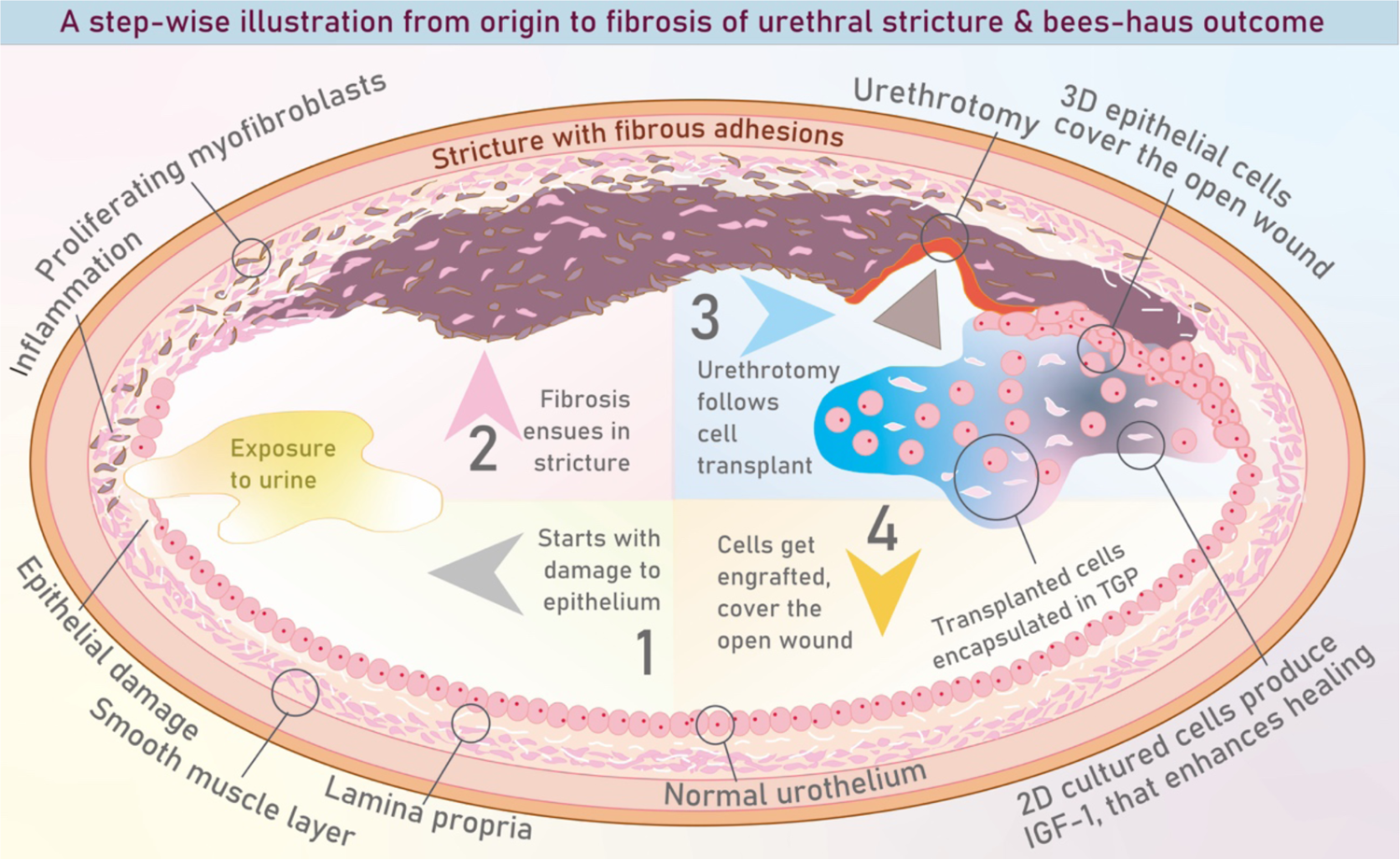

Illustration of pathogenesis of urethral stricture and the contribution of the hybrid combination of two-dimensional (2D) and three dimensional, 3D-TGP (Thermo-reversible gelation polymer) cultured cells to the successful repair of the stricture-inflicted male urethral epithelium in the buccal epithelium expanded and encapsulated in scaffold-hybrid approach to urethral stricture (BEES-HAUS) technique.

## Introduction

Urethral stricture (US) is defined as a constriction of the urethral lumen caused by ischemic spongiofibrosis. Trauma from urethral instrumentation, infection, and inflammatory illnesses have all been postulated as aetiologic reasons [1]. These cause the epithelial damage, which heals through fibrosis, resulting in a decrease of urethral lumen size and impaired urine flow and jeopardizes the reproductive function as well [2]. There are several surgical treatments for treating US, including urethral dilatation, internal urethrotomy, and urethroplasty. Anti-fibrotic medications and growth hormones have also been used to treat US. However, these surgical alternatives and adjuvant therapies might lead to additional problems, and there is recurrence of the stricture. As a result, existing treatment techniques for US can be considered to be of limited therapeutic effect [1].

Aside from pharmacological and surgical therapies, cellular and non-cellular regenerative medicine therapies have lately been tried. Clinical trial of submucosal injection of platelet-rich plasma (PRP) after DVIU [3] has been reported to reduce the recurrence of stricture in patients with primary, short bulbar strictures. Studies have also been conducted to investigate the efficiency of mesenchymal stem cells (MSCs) in the treatment of US [1]. Tissue engineering procedures combine a framework built of synthetic or natural biomaterials with cells and bioactive substances included into these constructions to activate the desired physiological response. Tissue engineering approaches for US have also been reported clinically including the autologous tissue-engineered buccal mucosa (TEBM) [4] in which keratinocytes and fibroblasts were isolated from buccal mucosa and cultured, seeded onto sterilised donor de-epidermised dermis and transplanted. There are regulatory approved tissue engineering products in some European countries [5] as well. Minced buccal mucosa suspended in fibrin gel termed as liquid buccal mucosa graft (LBMG), applied to the site of urethrotomy after DVIU has also been reported in animal models [6]. We have earlier reported a novel cell-based endoscopic approach called the buccal epithelium expanded and encapsulated in scaffold-hybrid approach to urethral stricture (BEES-HAUS) employing expanded and encapsulated buccal epithelial cells in a thermoreversible gelation polymer (TGP) scaffold in a clinical study in six patients [7]. After the procedure, all of the patients were able to void normally, with a peak flow rate of 24 mL/s on average. A 6-month urethroscopy revealed healthy mucosa at the urethrotomy site. In a subsequent animal study in rabbits, we demonstrated the proof of engraftment of transplanted buccal mucosal cells in BEES-HAUS over the urothelium at the urethrotomy site by immunohistochemistry markers positive for buccal epithelium, while negative for urothelium [8]. In this BEES-HAUS procedure, we utilized a combination of two-dimensional (2D) monolayer cultured cells and the three-dimensional (3D) TGP cultured cells that were transplanted using the TGP as a carrier [7–9] (Figure 1).

**Figure 1:**
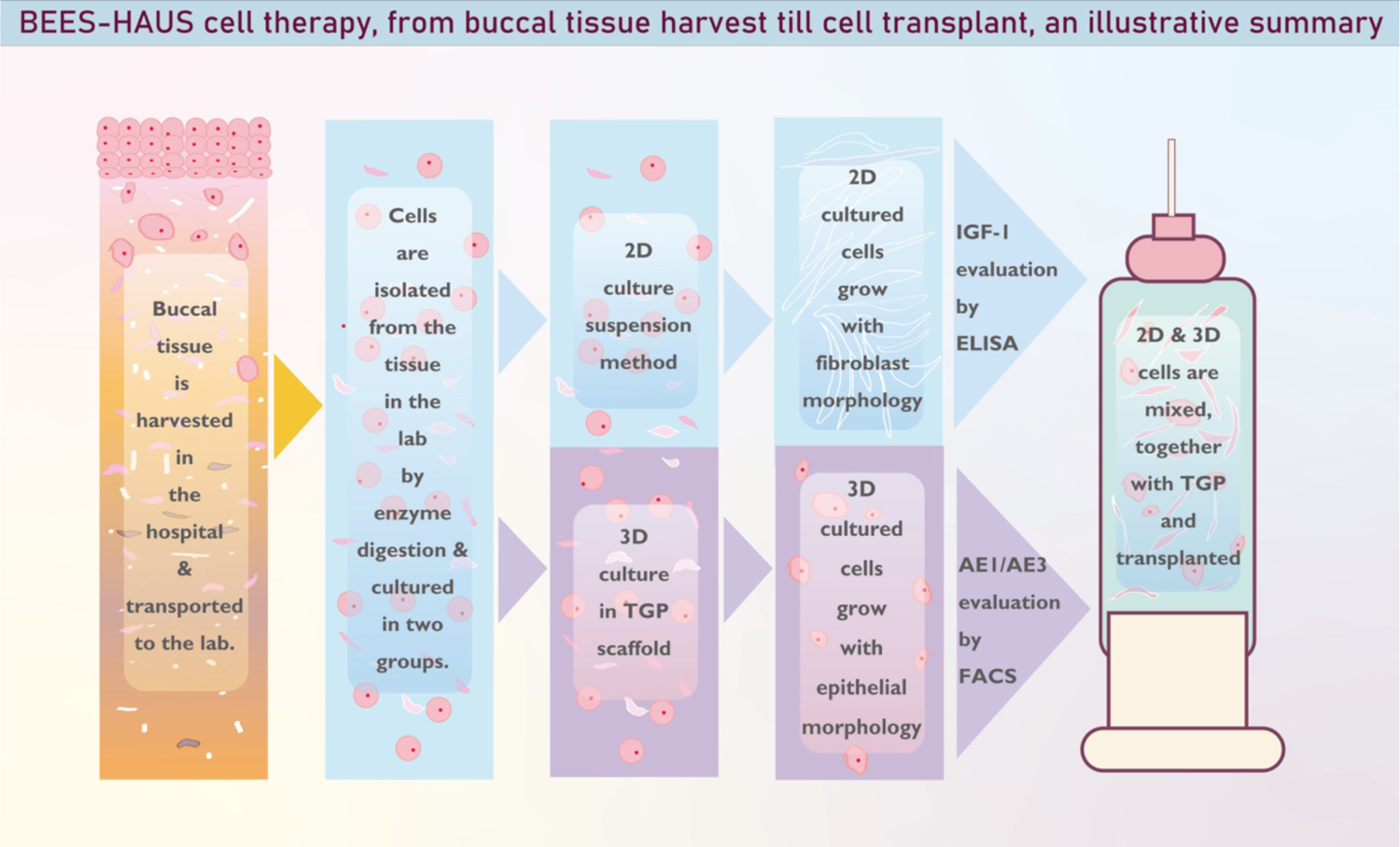
Buccal epithelium expanded and encapsulated in scaffold-hybrid approach to urethral stricture (BEES-HAUS) technique in which a combination of 2D monolayer cultured cells and 3D-TGP cultured cells are combined and transplanted, using the TGP as a carrier.

In the current study we have evaluated the advantages of such a cells’ combination that were employed for transplantation along with the 3D-TGP scaffold in the BEES-HAUS technique [7,8], by evaluating for IGF-1 secretion and AE1/AE3 pancytokeratin marker in the cultured cells before utilizing them for transplantation, thereby technically documenting their contribution to the successful outcome.

## Methods

The study was done in accordance with the declaration of Helsinki, following all prevailing guidelines and regulations. The study was approved by the ethical committee of National Defence Medical College, Japan (Ethics Committee Approval number: 4154 (9 April, 2020). Twenty two (n=22) human buccal tissue samples were obtained from adult patients undergoing biopsy for buccal mucosal graft urethroplasty, which are redundant after use in surgery. The tissues were transported in TGP provided by M/s GN Corporation in Japan. For use in transportation and culture, TGP was reconstituted with 10 ml of Dulbecco’s modified Eagle’s medium (DMEM)/F12 medium (Gibco BRL, Gaithersburg, MD, USA). Furthermore, this reconstituted TGP was stored at 4°C until usage. The tissues were subjected to enzymatic digestion using 1 ml of Digestion medium consisting of 1,000 PU/ml Dispase I (Oenon, Japan) in Dulbecco’s modified Eagle’s medium (DMEM) (Invitrogen) at 37°C overnight and then the epithelial layer was peeled off, minced and subjected to 0.5 ml of Accutase (Sigma) digestion fo 15 min at 37°C. The digested cells were washed twice at 1500 rpm for 10 min. These cells were counted with the trypan Blue dye exclusion method. For each sample, the cells obtained were divided and equally seeded in two groups. For group I (2D), the cells were seeded in 12-well Tissue Culture (TC) Plates (Greiner Bio-one, Austria) and overlaid with culture medium with 10% autologous serum. For group II (3D-TGP), the cells were mixed with reconstituted cold liquified TGP and added to 12-well TC plates in cold conditions. The TGP with the cells was allowed to solidify for 1 min. After the TGP was solidified, culture medium with 10% autologous serum was added over the TGP. In both the groups, the culture medium comprised of DMEM with 1% penicillin/streptomycin (P/S), 50 µg/mL gentamycin (GM), 0.25 µg/mL Amphotericin B (Amp) B, 5 µg/mL human insulin (91077C, Sigma, USA), and 10 ng/mL human epidermal growth factor (hEGF). Both the groups were cultured in 5% CO2 at 37°C for a maximum of 28 days. After culture of 28 days, the cells from 2D and 3D-TGP were harvested as per Horiguchi et al [7,8] and Katoh et al [9] and subjected to histological staining, flow-cytometry for AE1/AE3, CD140b and the supernatant from the cultures were subjected to ELISA for IGF-1 evaluation.

For histological staining, H & E staining was done as per standard protocol. For flow-cytometry, the cells were centrifuged and stained with the following isotype antibodies and antigen-specific antibodies. The cell samples were acquired on a FACSVia cell analyzer (BD Biosciences) and analyzed using FlowJo software (BD Bioscience)

### Antibodies list

① PE-AE1/AE3 (NSJ Bioreagents, cat# V2330PE-100T)
② FITC-CD31 (BioLegend, cat# 303103)
③ PerCP/Cy5.5-CD326 (BioLegend, cat# 369803)
④ APC-CD140b (BioLegend, cat# 323608)

(Isotype control antibodies were purchased from BioLegend and BD.)

For ELISA of the supernatant, Human IGF-I/IGF-1 Quantikine ELISA Kit (R&D systems, Cat#DG100B) was employed.

### Statistical analysis

Microsoft Office Excel statistics package and GraphPad Prism were used for statistical analysis. T-test and Wilcoxon rank sum test was used for comparison between the groups. Statistical significance was set at p<0.05, and values were represented as mean ± Standard deviation (SD)

## Results

The initial cell count (mean ± SD)was 0.015 ± 0.01 × 10^6 cells. The final cell count in 2D culture was 0.095 ± 0.092 × 10^6 cells. The final cell count in 3D-TGP culture was 0.18 ± 0.12 × 10^6 cells. The difference in the cell count between 2D and 3D-TGP was statistically significant (p-value=0.02).

The cells in 3D-TGP grew well with epithelial morphology, while the cells in 2D grew slowly with fibroblast morphology (Figure 2).

**Figure 2:**
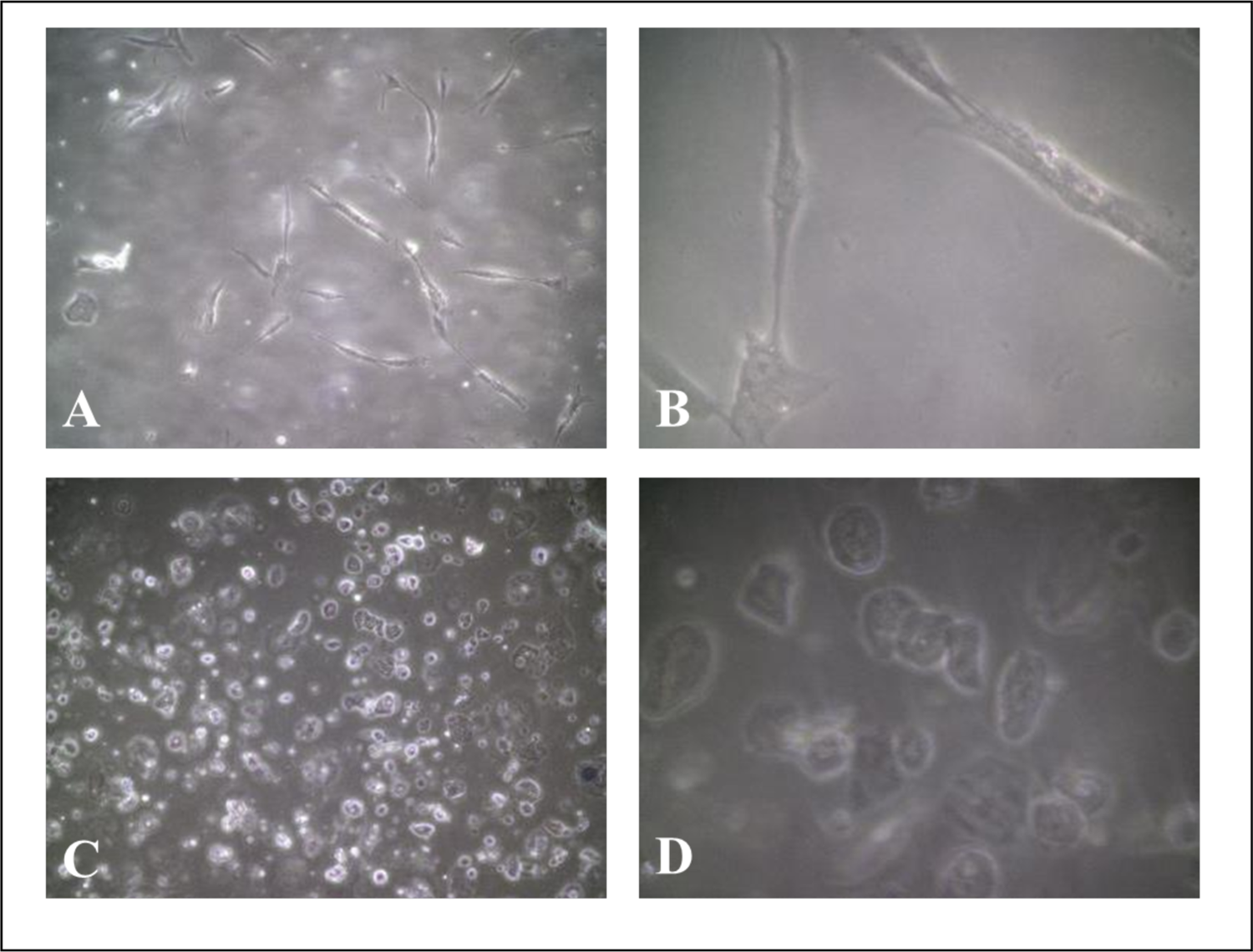
Culture images showed that 2D cultures had more cells with fibroblast morphology, A (x10 magnification) and B (x40 magnification) while 3D-TGP cultured images showed native epithelial morphology, C (x10 magnification) and D (x40 magnification

In H & E staining, the cells in 2D culture were intact as cells while there was a continuous tissue like growth in 3D-TGP culture (Figure 3).

**Figure 3:**
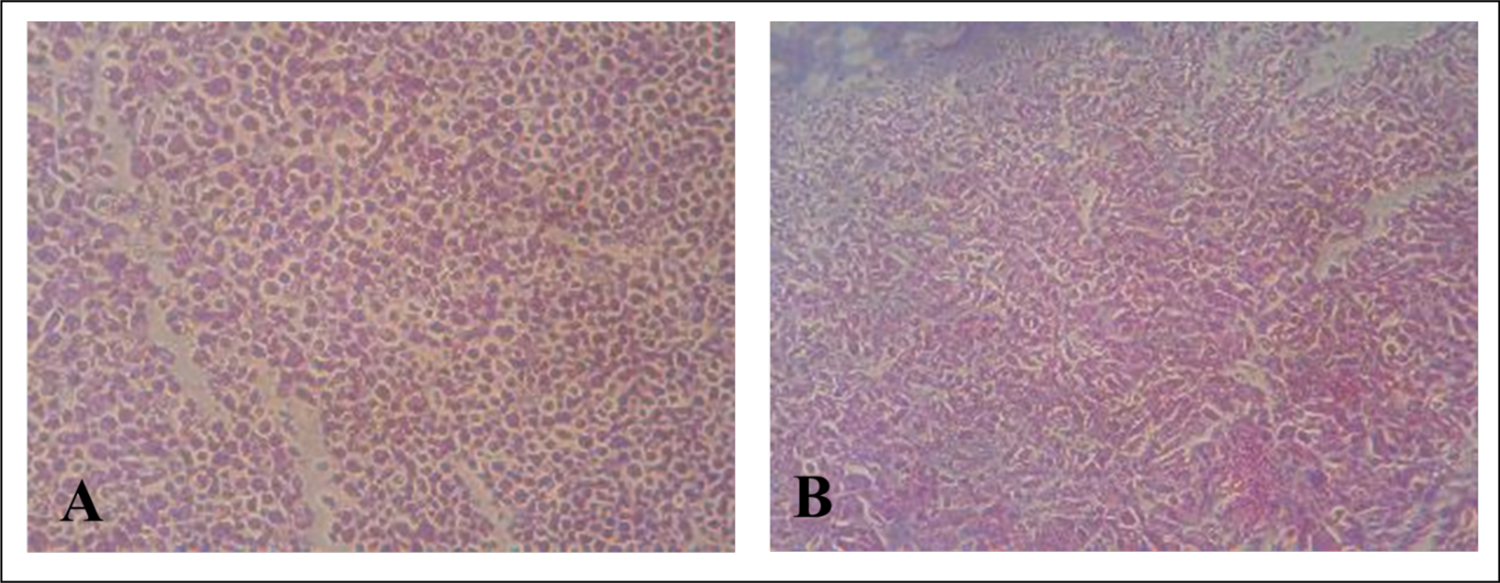
H and E staining showed that the cells in A. 2D were individual cells while B. 3D-TGP had a tissue like morphology

In flow cytometry, AE1/AE3 was negative on day 0 of cell culture and it became progressively positive in 3D-TGP cultures while the cells in 2D cultures were AE1/AE3 negative but CD140b positive indicative of fibroblast phenotype (Figure 4). The AE1/AE3 percentage after 28 days of culture was 1.05 ± 0.019 % in 2D while it was higher with statistical significance in 3D-TGP, 12.81 ± 0.063 % (p value < .00001) (Figure 5).

**Figure 4:**
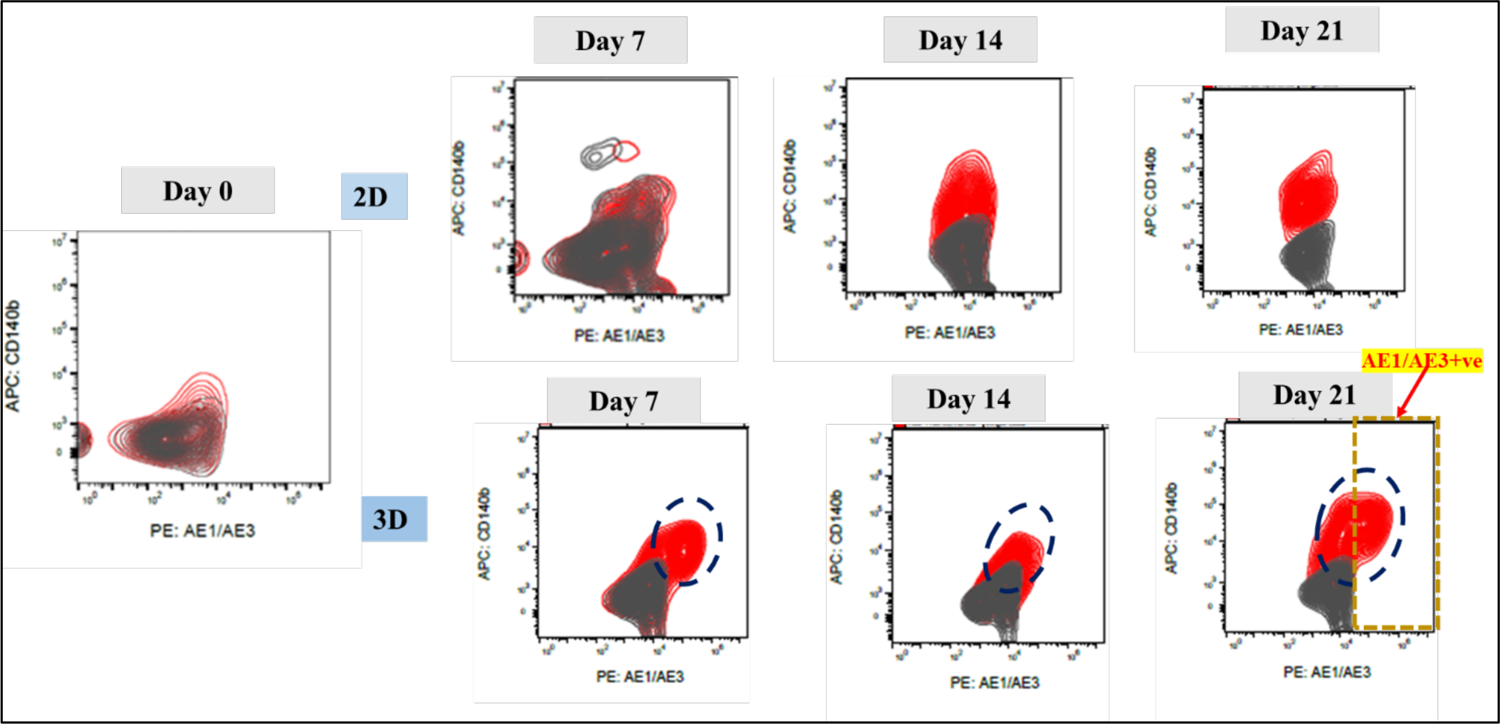
Flow cytometry images showing negative AE1/AE3 on day 0 of cell culture becoming progressively positive in 3D-TGP cultures while the cells in 2D cultures were AE1/AE3 negative but CD140b positive indicative of fibroblast phenotype

**Figure 5:**
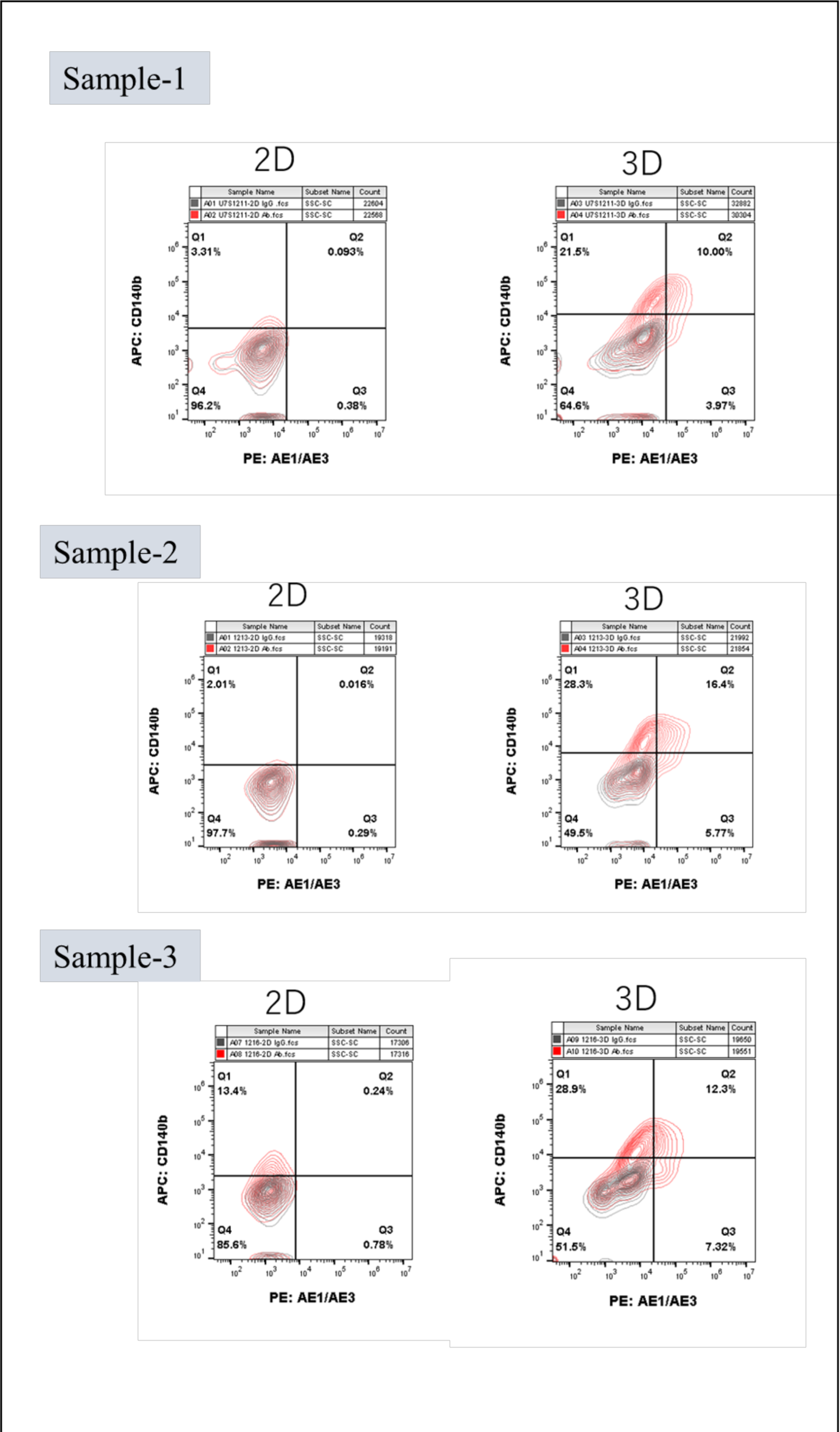
Flow cytometry images of three representative samples showing AE1/AE3 positive in 3D-TGP cultures while negative with CD140b positive in 2D cultures indicative of fibroblast phenotype

In ELISA of the supernatant for IGF-1, the level in 2D-TGP culture was higher 4.66 ± 1.50 ng/ml while in 3D-TGP it was 3.12 ± 1.20 ng/ml and the difference was statistically significant (p-value= 0.0002) (Figure 6)

**Figure 6:**
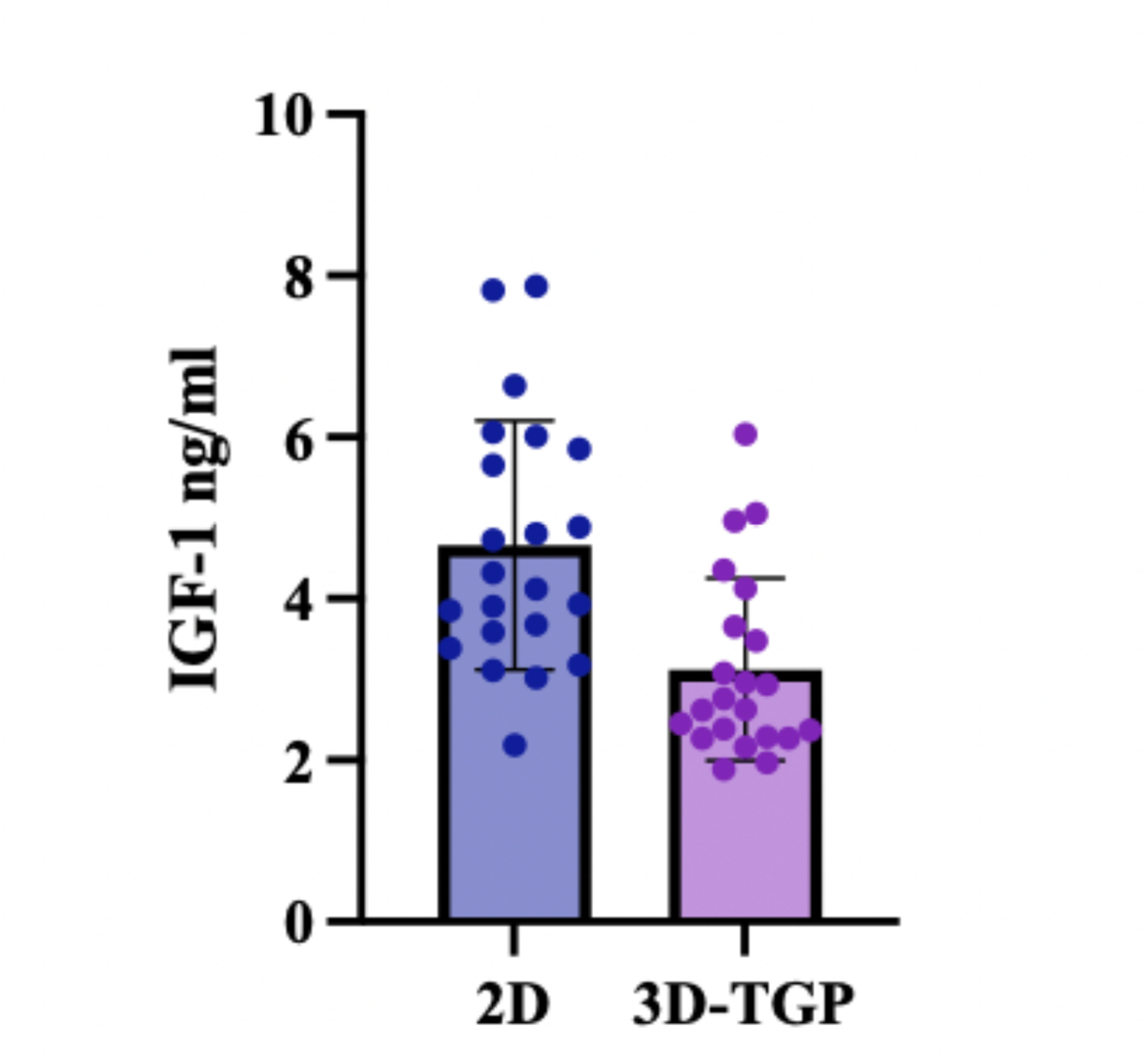
IGF-1 in supernatant of 2D cultures higher than 3D-TGP cultures

## Discussion

The BEES-HAUS method which has been proven to be efficacious in a human pilot study is one of the first of its kind report in the literature wherein, autologous buccal mucosal derived cells which were cultured in two different methodologies (2D-monolayer and 3D-TGP) together, as described in the present study have yielded a successful clinical outcome, following their transplantation using the TGP as a carrier into the urethral lumen, after releasing the fibrous strands causing the adhesion and stricture. The outcome is considered successful in terms of epithelialization of the stricture-inflicted area, previously covered with fibrous adhesions and the long term stricture recurrence-free patency, enabling proper voiding in human patients as well as a successful engraftment of the transplanted buccal tissue derived cells in animal models, that were culture expanded in the same manner as in human studies conducted [7–9]. Regenerative and tissue engineering cell based therapies contribute to repair the injured tissue’s function by either trans-differentiation to the cell type of interest in the transplanted host tissue, or manipulate the cell and molecular pathways regulating healing in order to replace damaged tissue and facilitate repair [10]. Growing evidence indicates that cell-mediated repair occurs as a result of paracrine substances being released into the surrounding tissue, which then control a range of healing processes, such as tissue, neovascularization, remodelling, and differentiation. Therefore, based on this concept of paracrine based regeneration, growth factors are applied for helping in regeneration. Earlier, Insulin-like growth factor-1 (IGF-1) has been linked to wound healing by increasing keratinocyte motility and promoting proliferative response in the wound. IGF-1 binds to its receptor which has been reported in rat urethra and has been shown to promote urothelial cell proliferation resulting in improved urethral wound healing via stricture prevention [11]. Shinichi et al have reported a study in which an insulin-like growth factor 1 sustained-release collagen urethral catheter significantly improved wound healing and prevented urethral stricture after urethral injury [12]. We have already reported the success of the BEES-HAUS procedure in clinical and animal studies of urethral stricture [7,8]. The rationale to use a combination of 2D and 3D-TGP cells for transplantation was to utilize the maximum quantity of cells that can be cultured in vitro as increased cell counts have been reported to increase the success of engraftment and repair [13]. Another rationale is that we don’t employ any feeder layers for culture but rather the TGP provides a 3D environment that promotes culture of cells preserving their native phenotype characteristics [14] and therefore combining cells from 2D would be an added advantage as it may secrete a few factors that would aid in regeneration after transplantation. Indeed, in the present study, the IGF-1 which is secreted more in 2D than 3D-TGP will add to the success of the BEES-HAUS procedure as it enhances wound healing and will also prevent further stricture recurrence. The fibroblasts in 2D will serve as a warehouse of other growth factors, as fibroblasts have been reported to secrete several growth factors and interleukins, such as interleukins 1, 6, and 8, granulocyte macrophage colony stimulating factor, transforming growth factor-α and -β, platelet-derived growth factor as well as members of the fibroblast growth factor family such as Keratinocyte growth factor (KGF), epidermal growth factor (EGF) etc [15]. The increased pancytokeratin AE1/AE3 expression in the 3D-TGP cells is a proof of their epithelial morphology maintenance throughout the culture period while the increased Cd140b in 2D cultured cells demonstrate their mesenchymal lineage conversion [16]. Since, epithelial morphogenesis, homeostasis, and repair are assumed to be dependent on epithelial-mesenchymal interactions and the mesenchyme has been hypothesized to promote epithelial development and differentiation by providing an appropriate biomatrix environment or by synthesising diffusible substances [17], this hybrid combination of 2D and 3D-TGP cultured cells for transplantation in BEES-HAUS serves as a successful technique for optimal urethral stricture repair and regeneration. Further validation of the underlying molecular mechanisms and studying the applicability of BEES-HAUS in other epithelial regeneration such as oesophageal stricture treatment, intestinal mucosa repair etc., is warranted in further studies. A significant contribution of this study to regenerative therapies is the use of the chemically synthesized TGP scaffold which has been earlier reported to support the in vitro culture of different kinds of cells and stem cells such as corneal limbal epithelial stem cells [18], corneal endothelial precursor cells [19], bone marrow mononuclear cells [20], hepatocytes [21], retinal pigment epithelial cells [22], chondrocytes [23] and pluripotent stem cells [24] apart from helping in transportation of cells and tissues without the need for cool preservation [25, 26] and serving as a transplantation scaffold for cell and tissue constructs in pre-clinical and clinical studies including limbal stem cell deficiency (LSCD) [27], spinal cord injury [20], alveolar bone regeneration [28] and urethral stricture [7–9]. In this study, we would like to emphasize that the 3D-TGP when used as scaffold for transplantation in the BEES-HAUS technique [7–9] has allowed the migration of cells within it, that could have aided the engraftment of cells in humans and rabbits which also has been documented in corneal limbal stem cell transplantation in an animal ocular surface model [29]. This TGP polymer also has provided an in vitro platform which allows the growth of 3D cells but since it doesn’t allow fibroblasts to grow, wherein a 2D culture system is needed in vitro, which under in vivo conditions becomes an advantage, as TGP doesn’t allow fibroblasts to grow and may have played a contributing role in preventing fibrosis in urethral stricture, which needs further research.

One of the limitations of this study was the inability to measure the exact hostile nature of the in vivo milieu in both human and animal studies because there is no investigation system presently available that allows the nature and severity of the pathological nature of the fibrotic area exactly, especially in the human urethral stricture and also the dynamically evolving healing process during which several cytokines and inflammatory and parameters keep interacting to bring back a homeostasis. The hybrid combination of cells transplanted have yielded a repair towards normalcy; however apart from the IGF-1 several other parameters from the transplanted cells could also have played a role in this, which needs further analysis, for fine tuning the application. Second limitation is the quantity of cells to be transplanted, which unless the previous question on the quantum of the pathological involvement is known, would be difficult to arrive at. Third is the difference between the animal models and the human disease in terms of chronicity in human pathology and multiple interventions during the repetitive interventions further making the already hostile tissue environment worse, against an animal study where the damage is made only once, that too in a young animal. Another important point to be considered is the other specific applications where similar buccal tissue derived cells have been applied as buccal mucosal epithelium has also been tried for regeneration of the cornea in which they have been able to replace the transparent corneal layer but the in vitro culture in that methodology uses feeder layers [30, 31]. From this viewpoint, this study having been proven successful engraftment in vivo earlier [7–9], may have several unknown inputs similar to the above mentioned corneal regeneration wherein a 3T3 feeder layer provided the necessary paracrine inputs and what we have accomplished as successful regeneration in vivo remains to be studied under in vitro environments, which may yield more insights into the interaction of such transplanted hybrid cells of different nature with the in vivo factors. These insights to be gained may continue to remain a limitation until we develop 3D organoid systems that can exactly recapitulate the dynamically changing in vivo tissues, under in vitro conditions.

## Conclusion

The present study has demonstrated that a hybrid combination of 2D and 3D-TGP cultured buccal mucosal cells complement each other in successfully repairing a stricture-inflicted urethral epithelium as proven in the BEES-HAUS technique, as the 3D-TGP cells contribute to the epithelial regeneration based on higher AE1/AE3 expression while the 2D cultured cells of fibroblast morphology secreted IGF-1, aiding the healing and together preventing recurrence, when transplanted using a polymer scaffold (TGP) as a carrier which ably and passively supported the engraftment of the cells. Further investigations of these kind of hybrid culture systems of different environments and cells may not only pave way for regenerative therapies for epithelial regeneration, but also other types of tissues based on their nature and the environment where damaged or dysfunctional cells or tissues have to be repaired or restored or replaced or regenerated.

## Acknowledgements

The authors thank

1. Mr. Mathaiyan Rajmohan, Mr. Ramalingam Karthick of Nichi-In Centre for Regenerative Medicine (NCRM) for their assistance with the cell culture work described in the manuscript.
2. Ms. Yoshiko Amikura and staff of GN Corporation Co Ltd, Japan for their liaison assistance with the conduct of the study.
3. Mr. Hirotaka Morito of JBM Inc., Dr. Fumihiro Ijima and Dr. Hiroshi Hirano of Hasumi International Research Foundation, Tokyo, for their technical assistance with the data collection.
4. Ms Eiko Amemiya of II Dept of Surgery, Yamanashi University for her secretarial assistance.

## Notes

### Competing Interest Statement

Author Dr Katoh is an inventor to several patents of relevant methods and products described in this manuscript and a member of the governing board of Jinsei-sha, a social welfare trust, that owns the Edogawa Hospital; Author Dr. Abraham is an inventor of several patents on biomaterials and methodologies including the one described in the manuscript and a shareholder in GN Corporation, an applicant to IP rights.

